# IL-36/IL-36R Signaling Promotes CD4+ T Cell-Dependent Colitis via Pro-Inflammatory Cytokine Production

**DOI:** 10.1101/2023.05.24.542162

**Authors:** Maya Maarouf, Michal Kuczma, Timothy L. Denning

## Abstract

Inflammatory bowel disease (IBD) is a multifactorial, chronic disease that affects approximately 1.5 million people in the United States [1]. It presents with inflammation of the intestine with unknown etiology and its two main forms are Crohn’s disease (CD) and ulcerative colitis (UC). Several important factors are implicated in the pathogenesis of IBD, one being dysregulation of the immune system resulting in the accumulation and stimulation of innate and adaptive immune cells and subsequent release of soluble factors, including pro-inflammatory cytokines. One of these cytokines is a member of the IL-36 cytokine family, IL-36γ, which is overexpressed in human IBD and experimental mouse models of colitis. In this study, we explored the role of IL-36γ in promoting CD4^+^ T cell activation and cytokine secretion. We found that IL-36γ stimulation of naïve CD4^+^ T cells significantly induced IFNγ expression *in vitro* and was associated with augmented intestinal inflammation *in vivo* using naive CD4^+^ cell transfer model of colitis. Using IFNγ-/- CD4^+^ cells, we observed a dramatic decrease in the ability of TNFα production and delayed colitis. This data not only suggests that IL-36γ is a master regulator of a pro-inflammatory cytokine network involving IFNγ and TNFα, but also highlights the importance of targeting IL-36γ and IFNγ as therapeutic approaches. Our studies have broad implications in relation to targeting specific cytokines in human IBD.

## Introduction

Inflammatory bowel disease (IBD) is a multifactorial, chronic condition that remains incurable until this day. Its two main forms are Crohn’s disease (CD) which affects any part of the GI tract, and ulcerative colitis (UC) which impacts the colon only [2]. Symptoms of IBD include abdominal pain, loss of appetite, diarrhea, and weight loss [2]. To date, the therapeutics available currently primarily work to reduce and suppress inflammation, without curing the disease [3]. Unfortunately, the etiology of IBD remains ambiguous. Several factors play a role in the pathogenesis of IBD, including genetics, the microbiota, environmental factors, and immunological dysregulation [4,5].

A disruption in the mucosal barrier increases intestinal permeability [6,7], resulting in immune dysregulation with an extensive infiltration of innate and adaptive immune cells. These events lead to intestinal inflammation and IBD. The intestinal epithelial barrier acts as a defense barrier between the intestinal mucosa and the lumen, composed of a layer of intestinal epithelial cells (IECs) [8]. During a primary immune response, the priming of T cells occurs in the mesenteric lymph nodes (mLN), where naïve CD4^+^ T cells interact with the antigen and differentiate into activated effector cells. These effectors can polarize into any of the different subsets, Th1, Th2, Th17, Th22 or T regulatory cells, depending on the microbial challenge to be eliminated [9,10]. Th1 cells, especially present in CD patients, express the transcription factor T-bet and function by producing IFNγ and TNFα. On the other hand, UC is associated with Th2-mediated responses, including elevated production of IL-4, IL-5, and IL-13. Th17 cells are also found in IBD patients, and this subset plays a role in eliminating extracellular pathogens and maintaining commensal microbiota by primarily producing IL-17A and IL-17F [10]. Th17 differentiation is induced by IL-6 and RORγT. Another IBD-associated Th subset are Th22 cells. These cells are known for their production of IL-22 and expression of RORγT and are negatively regulated by T-bet. All the mentioned subsets and cytokines are classified as pro-inflammatory. Regulatory T cells (T regs) produce anti-inflammatory cytokines. They are known for their inhibitory effects and ability to dampen immune responses. These cells express Foxp3 and function by producing IL-10 [7]. While CD4^+^ T cells are strongly associated with IBD pathogenesis [11], it is the cytokine imbalance of pro- vs. anti-inflammatory factors dictating the fate of Th lineage commitment [12].

The IL-1 superfamily of cytokines is important in innate and adaptive immunity. This superfamily consists of the following cytokines: IL-1α, IL-1β, IL-18, IL-33, and IL-36. IL-36 includes three agonists: IL-36α, IL-36β, and IL-36γ. These ligands and their receptor IL-36R are expressed mostly on epithelial cells, and some of their target cells include intestinal epithelial cells and naïve CD4^+^ T cells [13,14]. Binding to their receptor, IL-36R, activates nuclear factor kappa B (NF-κB) and mitogen-activated protein kinases (MAPK) [13], causing inflammation. However, IL-36 cytokines are regulated by their natural inhibitor, the antagonist IL-36Ra, which inhibits the inflammatory signaling pathway [15]. In IBD patients, IL-36α and IL36γ levels were found to be elevated [16]. IL-36 signaling promotes inflammation, making IL-36/IL-36R potential therapeutic targets for IBD. Common ways to block IL-36R signaling include anti-IL-36R antibodies and receptor antagonists. Studies using the anti-IL-36R antibody led to the development of drugs like Spesolimab (a humanized monoclonal IgG1 antibody) [17]. This receptor blocking antibody prevents all IL-36 ligands from binding to the receptor [18]. Spesolimab showed clinical efficacy in patients who have the autoimmune disease, psoriasis, in phase I studies. It is currently being tested in CD patients and is in phase II trials in active UC patients [17].

While significant progress has been made towards managing IBD, understanding the role of IL-36 cytokine family in IBD pathogenesis is crucial, since the antagonist IL-36γ is elevated in human IBD and experimental colitis, as well as other inflammatory diseases. While anti-TNFα therapy can induce clinical remission in CD patients [19], the disease ends up relapsing in more than 60% of patients [20]. Cytokine blockers and immunomodulators were also recently developed to treat intestinal inflammation [21]. However, patients can still go into clinical relapse. Therefore, evaluation of IL-36/IL-36R signaling in T cell-mediated colitis will further narrow down therapeutic targets for intestinal inflammation.

## Materials and Methods

### Animal Models

The following mice were obtained from the Jackson Laboratory: Wild-type C57BL/6 (B6 WT), Rag-/-, and B6.129S7-*Ifng*^*tm1Ts*^/J (IFNγ-/-). All animal procedures were performed according to the GSU (IACUC) guidelines. All mice involved in experimental procedures were eight to twelve weeks old, both females and males.

### Cell Culture, RNA Isolation for QPCR, and Supernatant Harvesting for ELISA

Spleens and lymph nodes were collected from wild-type and IFNγ -/- mice and processed into single-cell suspensions by passing through 40μm mesh (Fisher). Splenocytes were resuspended in Ammonium-Chloride-Potassium (ACK) buffer (Thermo) for five minutes to lyse erythrocytes then washed once with PBS. Cells from splenocytes and lymph nodes were combined to isolate naïve CD4^+^ T cells (CD4^+^CD25^-^CD45RB^+^) via EasySep™ Mouse Naïve CD4^+^ T Cell Isolation Kit (Stemcell Technologies) and the purity was assessed with FACS by labeling cells with antibodies specific to mouse CD4 (clone GK1.5), CR45RB (clone MB4B4) and CD25 (clone PC61).

For cell culture, 96-well flat bottom plates were coated with 5 μg/mL αCD3 (clone 145-2C11) and 1 μg/mL αCD28 (clone 37.51). Naïve CD4^+^ T cells were plated at 2×10^5^ cells per well, in the presence or absence of IL-36α, IL-36β, and IL-36γ IL-36γ (R&D Systems) at a concentration of 100 ng/mL and cultured for 48 hours in the TC incubator (37°C, 5% CO_2_). In some experiments, αIFNγ neutralizing antibody (clone XMG.1, 2 μg/mL) (Thermofisher) was added to a cell culture. At the indicated time, supernatants were collected for IFNγ ELISA done following the manual included in the kit (Invitrogen) and kept at -80°C until use. RNA was isolated with RNeasy Mini kit (Qiagen) according to manufacturer’s instructions and DNA-free RNA was reverse-transcribed with Superscript IV kit per supplier’s protocol (Invitrogen).

### Adoptive T Cell Transfer Colitis Model

Splenocytes and lymph node cells were harvested from eight-week-old wild-type and IFNγ-/- mice. Naïve CD4+ cells were enriched as described above, resuspended at 2.5×10^6^ cells/ml in sterile PBS and 200 μl (0.5×10^6^ cells) were i.p injected into Rag-/- mice. Mice were followed for a period of 10-12 weeks and sacrificed when they lost 20% initial body weight.

### Hematoxylin and Eosin (H&E) Staining and Histological Score Evaluation

Colons were flushed from feces and fixed with 10% neutral buffered formalin (Fisher) for 72 hours, moved to 70% ethanol and submitted for processing and H&E staining to HistoWiz (Brooklyn, NY). Histological scoring was performed in a blind matter, using a 0-4 scale (0, no discernible inflammation; 1, small, focus of inflammation; 2, small, multiple foci of inflammation; 3, large focus of inflammation; and 4, large, multiple foci of inflammation).

### Quantification of Fecal Lipocalin (LCN2) by ELISA

Fecal pellets were collected from mice during the course of disease and frozen at -80°C until use. Pellets were resuspended at 100 mg/ml in PBS left at 4°C overnight, to allow the feces to soften. Samples were then homogenized for 1 minute, centrifuged (16,000*g*, 15 minutes, 4°C) and supernatant was aliquoted and frozen at -80°C until use. To analyze lipocalin (LCN2) levels, Mouse Lipocalin-2/NGAL DuoSet ELISA kit by R&D Systems was used following the manufacturer’s protocol.

### Intracellular Cytokine Staining (ICCS)

ICCS was done in cLP, mLN and spl cells. In order to isolate immune cells from colon tissue, it was cut longitudinally, feces were removed, and the tissue was extensively washed in PBS after which, 0.5 cm fragments were incubated in the heated shaker in HBSS buffer containing 10% FBS and 2mM EDTA at 37°C for 15 min with the mild shaking (150 rpm) to remove epithelium. Then, enzymatic digestion of tissue pieces was done with collagenase D (1 mg/ml) and DNAseI (0.1 mg/ml) (both from Roche) in HBSS/10%FBS for 45 min (37°C, 150 rpm). Finally, tissues were washed with HBSS, filtered through 70μm mesh and run through glass wool-packed columns to purify immune cells. Cell suspension was filtered through 40μm mesh and used for stimulation followed by FACS staining. Briefly, cells (cLP, mLN, spl) were plated in complete culture medium supplemented with 50 ng/mL PMA, 1 μg/mL ionomycin (both from Sigma), 1:1000 Brefeldin A, and 1:1000 monensin (both from BioLegend). Cells were stimulated for 4 hours in the TC incubator. After that time, cells were washed once with PBS and stained with fixable aqua viability dye (Invitrogen) for 15 min at RT. After washing, FcR was blocked with 2.4G2 Ab (BioXCell) and stained with antibodies specific to CD4, CD44, and Ly6G for 20 minutes in the dark. The cells were then washed with PBS, fixed in Fixation Solution (eBioscience) for 20-30 minutes in the dark at room temperature, permeabilized with Perm Solution (eBioscience) for 20 min at RT and stained with anti-IFNγ and anti-TNFα (clone MP6-XT22) antibodies for 30 min at RT. After washing, cells were resuspended in PBS for flow cytometer analysis.

### Statistical Analysis

All statistical analyses were performed with GraphPad Prism software, version 9.0 (Graphpad Software). ONE-way ANOVA and Tukey’s Multiple Comparison Test or Student’s t test were used to determine significance. *P < 0.05, **P < 0.01, ***P < 0.001; ****P <0.0001; ns= not significant.

## Results

### IL-36*γ* Increases Naïve CD4^+^ T Cell Proliferation and IFN*γ* Production *In Vitro*

Previous studies have shown that IL-36β can increase naïve CD4^+^ T cell proliferation and induce pro-inflammatory cytokine production and Th1 polarization [15]. We choose to investigate the influence of the three IL-36 ligands, IL-36α, IL-36β, and IL-36γ, on naïve CD4^+^ T cells. naïve CD4^+^ cells were stimulated with αCD3/αCD28 Abs for 48 hours in the presence or absence of any of the IL-36 ligands. As a negative control, some cells were left unstimulated. At 48 hours, cells were counted and cell number was significantly higher in cultures stimulated in the presence of IL-36γ compared to other conditions (Fig. 1A). We also observed more cell clusters under this condition (data not shown). To further assess cell activation, we checked for the activation marker by flow cytometry. The plots in Fig. 1B&C show a major increase in CD4^+^CD44^+^ expression on cells activated in the presence of IL-36γ. In addition to enhanced cell proliferation and activation, we observed significant induction of IFNγ in IL-36γ-stimulated cells (Fig. 1D), compared to the control sample (cells stimulated with αCD3/αCD28) and samples that received IL-36α or IL-36β.

**Figure 1.**
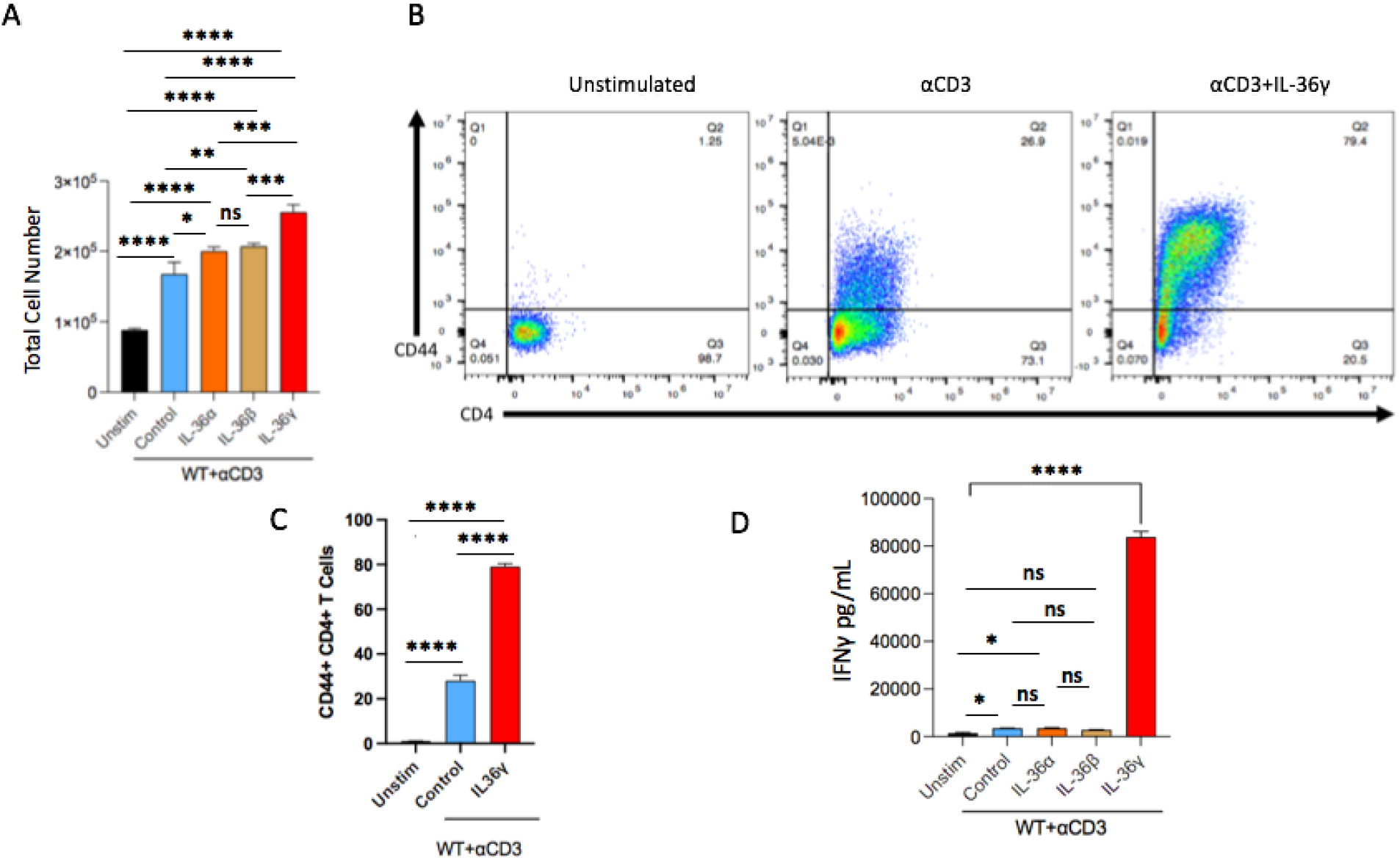
IL-36*γ* Increases Naïve CD4^+^ T Cell Proliferation and IFN*γ* Production *In Vitro*. (A) Naive CD4^+^ T cells were isolated from splenocytes and were stimulated on a αCD3/αCD28 pre-coated plate in the presence of IL-36α, IL-36β, or IL-36γ. Some cells were left unstimulated as a control. These data were collected in 48 hours. Cells were counted under a hemocytometer. (B&C) The expression of the activation marker CD44 was evaluated after 48 hours, which was found to be highly expressed on cells stimulated with IL-36γ. (D) Supernatants from these cultures were checked for the pro-inflammatory cytokine IFNγ by ELISA. All data are presented as mean ± SEM; *P < 0.05, **P < 0.01, ***P < 0.001; ****P<0.0001; ns= not significant, one-way ANOVA with Tukey’s multiple comparison test.

### Transfer of IL-36*γ* Stimulated Naïve CD4^+^ T Cells to Rag-/- Mice Exacerbates T Cell-Mediated Colitis

Next, we evaluated the impact of the robust IFNγ secretion by IL-36γ on T cell-mediated colitis. Cells activated *in vitro* in the presence or absence of IL-36γ, were transferred to Rag-/- mice. As a control, unstimulated naïve CD4^+^ T cells were transferred as reported [11]. Weight, mouse activity, stool consistency, and fecal LCN2 levels were monitored over the span of 10 weeks. The mice that were transferred with activated cells displayed much lower activity than the mice that received the unstimulated cells. Mice that received cells stimulated with IL-36γ cells also developed softer stool and secreted the highest fecal LCN2 levels, when checked by ELISA (Fig. 2A). At the time of sacrifice, the spleens and colons were collected for analysis. Colons from the IL-36γ group were shorter in length than the other two groups and appeared to be more thickened and inflamed. Moreover, it is obvious that the unstimulated group colons portray distinct fecal pellets, the stool progressed to a softer consistency in the other two groups, especially in the group that was treated with IL-36γ-stimulated cells (Fig. 2B). The latter also developed a visibly enlarged spleen (Fig. 2C). H&E staining showed that the same group had the most leukocyte infiltration (Fig. 2D), and therefore, the highest histological score (Fig. 2E).

**Figure 2.**
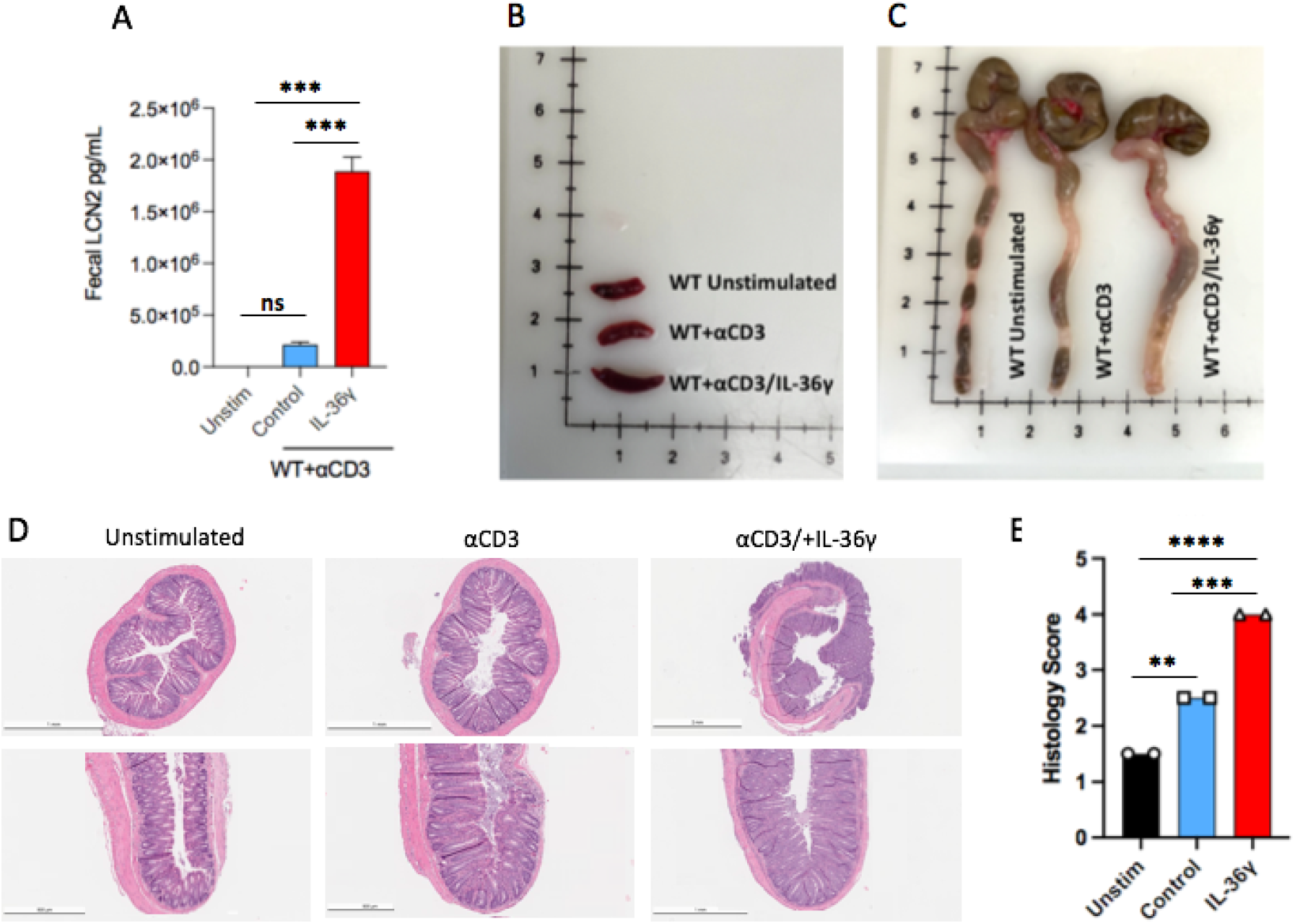
Transfer of IL-36*γ* Stimulated Naïve CD4^+^ T Cells to Rag-/- Mice Exacerbates T Cell-Mediated Colitis. (A) Lipocalin 2 (LCN2), was measured by ELISA from fecal samples at time of sacrifice from mice that were adoptively transferred with unstimulated naïve CD4^+^ T cells or naïve CD4^+^ T cells stimulated with αCD3 +/- IL-36γ. (B&C) Images of spleens and colons, show an enlarged spleen and a shorter colon length, along with thickened tissue in mice that received naïve CD4^+^ T cells stimulated with IL-36γ. (D) Representative H&E staining of colon sections from the mentioned mice groups is shown. (E) Histology scoring of colon sections from mice that were treated as mentioned. All data are presented as mean ± SEM; *P < 0.05, **P < 0.01, ***P < 0.001; ****P<0.0001; ns= not significant, one-way ANOVA with Tukey’s multiple comparison test.

### The absence of IFN*γ* Reduces TNF*α* Production by Naïve CD4^+^ T Cells

To test the link between IL-36γ and IFNγ, we carried out similar *in vitro* experiments on naive CD4^+^ T cells using an IFNγ neutralizing antibody (αIFNγ). The cells were stimulated with αCD3/αCD28 and IL-36γ, in the presence of αIFNγ or an isotype antibody. As a control, some cells were stimulated without the cytokine but received the isotype antibody. After 48 hours, the cells were counted. We observed a higher cell number in the samples that received IL-36γ, compared to the sample that only received the isotype antibody (Fig. 3A). Another pro-inflammatory cytokine, TNFα, was evaluated via ELISA. While IL-36γ did induce TNFα production by naive CD4^+^ T cells, neutralizing IFNγ reduced its secretion (Fig. 3B), which suggests that IFNγ and TNFα function together in a pathway, where IFNγ is upstream of that pathway. Not only that, but when IFNγ-/- naïve CD4^+^ T cells were transferred to Rag-/- mice to assess intestinal inflammation, there was a reduction of TNFα in the mesenteric lymph nodes that was assessed by intracellular cytokine staining (ICCS) (Fig. 3E).

**Figure 3.**
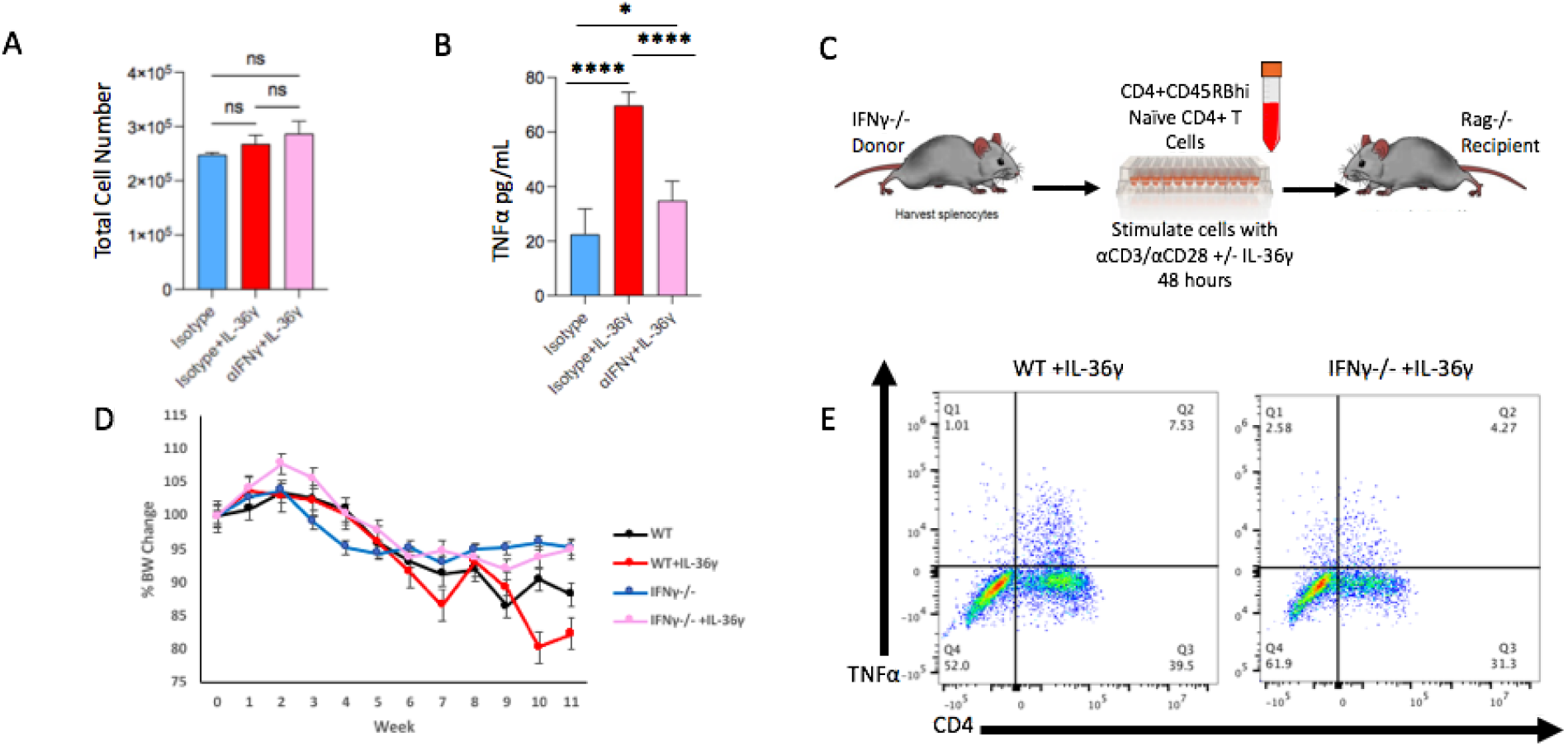
IFN*γ*-/- Naïve CD4^+^ T Cells Stimulated with IL-36*γ* Produce Less TNF*α*. (A) Cell number of naive CD4^+^ T cells stimulated with αCD3/αCD28 +IL-36γ, either with αIFNγ or an isotype antibody-at 48 hours. As a control, some cells were stimulated but only received the isotype. (B) TNFα concentrations evaluated by ELISA. (C) Diagram showing the adoptive transfer model using donor IFNγ-/- mice as donors and Rag-/- recipients. Naive CD4+ T cells are isolated and stimulated *in vitro* in the presence or absence of IL-36γ for 48 hours prior to the transfer. (D) Weight of the recipient mice was monitored during the time of the experiment over the course of 11 weeks. (E) After sacrifice, TNFα production in the mesenteric lymph nodes was evaluated by ICCS. All data are presented as mean ± SEM; *P < 0.05, **P < 0.01, ***P < 0.001; ****P <0.0001; ns= not significant, one-way ANOVA with Tukey’s multiple comparison test.

### IL-36*γ* Stimulated IFN*γ*-/- Naïve CD4^+^ T Cells Induce Less Robust Colitis

Next, we determined the influence of IFNγ on intestinal inflammation by transferring IFNγ-/- naïve CD4^+^ T cells to Rag-/- mice. Prior to the transfer, these cells were stimulated with αCD3/αCD28 *in vitro* in the presence or absence of IL-36γ (Fig. 3C). As a control, WT naïve CD4^+^ T cells were also stimulated under the same conditions. The mice were monitored over the span of 12 weeks as described above. Over time, it was obvious that mice transferred with WT cells +/- IL-36γ became less active and physically weaker, some even developing lethargy. On the other hand, the mice that received IFNγ-/- cells, in the presence or absence of IL-36γ, remained active. Mice that received WT+IL-36γ cells lost the most weight, followed by the WT group. On the other hand, mice that received the knockout cells had minimal weight loss (Fig. 3D). Stool samples were checked for LCN2 levels. WT+IL-36γ group had the highest LCN2 concentrations. Surprisingly, the group with the second highest LCN2 levels was IFNγ-/-+IL36γ. The other two groups, WT and IFNγ-/-, had mild fecal LCN2 (Fig. 4A). The colons of mice that received WT cells were noticeably shorter in length, especially WT+IL-36γ group, compared to the groups that received IFNγ-/- cells (Fig. 4B). WT+IL-36γ group also had a visibly enlarged spleen, while the spleens of the other three groups were fairly the same size (Fig. 4C). Histological staining showed the most thickening of the submucosa and leukocyte infiltration in WT+IL-36γ colons out of the four groups (Fig. 4D). However, IFNγ-/- +IL-36γ had more leukocyte infiltration than IFNγ-/-. The same colon tissue was stained for CD4 by immunohistochemistry staining, where both groups that received IL-36γ-stimulated cells had the most CD4 infiltration (Fig. 4E). In conclusion, mice that received WT+IL-36γ cells had the most intestinal inflammation due to having the shortest colon lengths, largest spleens, high leukocyte infiltration and LCN2 levels, and the highest histology score (Fig. 4F).

**Figure 4.**
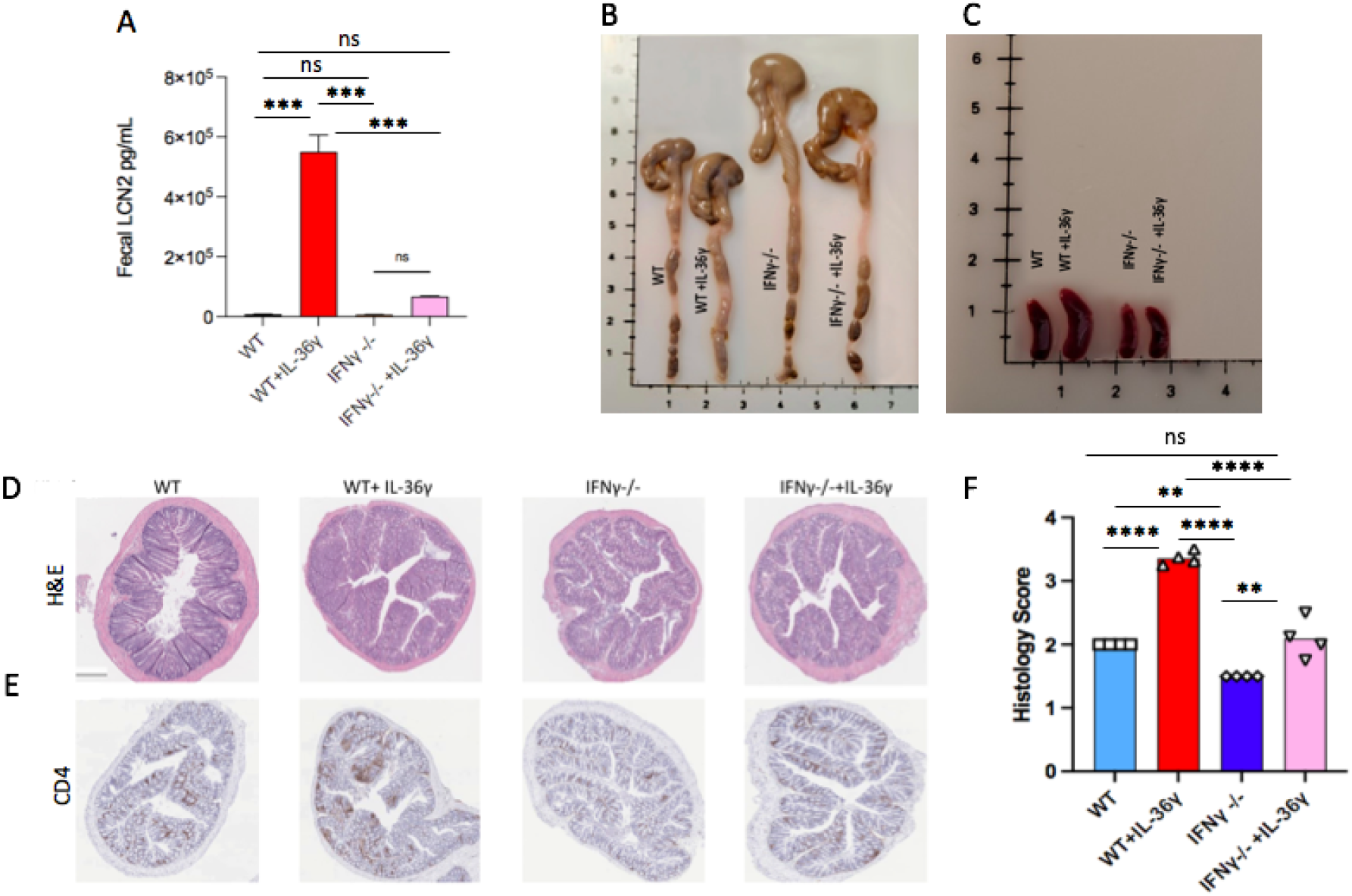
The Transfer of IFN*γ*-/- Naïve CD4^+^ T Cells Does Not Induce Robust Colitis. (A) Levels of fecal LCN2 levels were evaluated at time of sacrifice by ELISA. (B) Image of the harvested colons shows that mice that received cells stimulated with IL-36γ cause the most inflammation to the colon, making it the shortest in length. (C) Images of the harvested spleen show an enlarged spleen for the IL-36γ group. (D) H&E staining of colon sections from mice that received WT or IFNγ-/- naïve CD4^+^ T cells, in the presence or absence of IL-36γ. (E) Immunohistochemistry staining for CD4, of the same colonic tissue samples. (F) Histology scoring of colon sections from the same mice. All data are presented as mean ± SEM; *P < 0.05, **P < 0.01, ***P < 0.001; ****P <0.0001; ns= not significant, one-way ANOVA with Tukey’s multiple comparison test.

## Discussion

Cytokines are signaling molecules that play a crucial role in the pathogenesis of IBD. They are secreted by various immune cells in the gut to regulate different functions of the immune system. One cytokine that is observed to be elevated in IBD and experimental colitis is IL-36γ, which is a ligand in the IL-36 family. This cytokine is found to be elevated in other diseases where epithelial cells are impacted, such as skin condition psoriasis [9]. When it comes to IL-36γ signaling, some studies show a pathogenic role, while other studies found that IL-36γ can aid in healing of the disease. A study from our lab suggests a protective role for IL-36γ in the DSS colitis model by helping in barrier restoration [17,23]; however, another study from our lab suggests a pathogenic role for the cytokine in a more T cell mediated colitis model, using oxazolone [24]. Since IBD is heavily mediated by T cell responses, we investigated the influence of IL-36γ on naïve CD4^+^ T cells activated *in vitro* and in the adoptive transfer model which mimics the immunological mechanisms seen in human IBD [25].

When T cells are stimulated with antigens, they produce cytokines that can either be pro-inflammatory or anti-inflammatory. The cytokines secreted by T cells determine T cell differentiation. They can differentiate to Th1, Th2, Th17, or T reg cells, depending on the microbial challenge that needs to be eliminated. Two main subsets are observed during IBD, Th1 and Th17. During IBD, pro-inflammatory cytokines are produced in higher concentrations, compared to anti-inflammatory cytokines; therefore, exacerbating inflammation in the colon [26].

Our results demonstrate that stimulating naïve CD4^+^ T cells with αCD3/αCD28 Abs in the presence of IL-36γ significantly induces cell proliferation and expansion. IL-36γ-stimulated cells also produced high concentrations of IFNγ. We investigated the IL-36γ/IFNγ axis by transferring naïve CD4^+^ T cells stimulated with IL-36γ to immunocompromised Rag-/- mice. These mice lost weight over time, and their stools had a looser consistency, in addition to elevated levels of fecal LCN2. Their colons were shorter in length, had a thicker epithelium, higher histological scores, and more leukocyte infiltration. Since this suggests a critical for IFNγ in T cell mediated colitis, we continued the investigation by using naïve CD4^+^ T cells from IFNγ deficient mice, using the same method as WT mice. These cells were also stimulated *in vitro*, under similar conditions as WT cells, with IL-36γ. Another typical Th1 pro-inflammatory cytokine that we assessed was TNFα. TNFα production was suppressed when IFNγ was neutralized, suggesting a cytokine pathway that involves IFNγ and TNFα, in the presence of IL-36γ.

Rag-/- mice that received IFNγ deficient naïve CD4^+^ T cells, whether with or without IL-36γ stimulation, did not lose a significant amount of weight. Their activity remained healthy throughout the course of the experiment, with no signs of illness of lethargy. Lipocalin levels were assessed at time of sacrifice. WT+IL-36γ had the highest fecal LCN2 concentrations. IFNγ-/- +IL-36γ cell recipients showed a low induction of fecal LCN2 levels, but these levels were minimal.

## Conclusion

IL-36 cytokines promote the recruitment of immune cells, such as T cells [28]; and they induce the secretion of other pro-inflammatory cytokines, such as TNFα [29]. Our data showed that IL-36γ induced a notable expression and production of the Th1 pro-inflammatory cytokine IFNγ, by naïve CD4^+^ T cells compared with the other IL-36 family members confirming previous studies that showed IL-36 cytokines induce the production of IFNγ [30]. Moreover, our data suggests that blocking IFNγ can suppress the expression and secretion of the pro-inflammatory cytokine TNFα. Naïve CD4^+^ T cells stimulated with IL-36γ induced robust colitis upon transfer to Rag-/- mice, compared to control animals that received unstimulated naïve CD4 cells.

Based on the available pre-clinical studies on pro-inflammatory cytokines in the T cell mediated colitis model, understanding the role of IL-36γ in intestinal inflammation will highlight novel therapeutic approaches. In conclusion, our results demonstrate a potential for IL-36γ and IFNγ to be mutually targeted as therapeutics to alleviate colitis symptoms and perhaps prevent IBD, in the long run. Most of the available research targets downstream drivers of IBD, which narrows down the focus to specific molecules or pathways. Such downstream targets include TNFα, the IL-12/IL-23 axis, IL-23/IL-17 axis, JAK inhibitors, integrins and receptor modulators [31,32]. Some antibodies and biological therapies were developed to interfere with the function of these molecules and pathways, such as Anti-TNF therapy, anti-IL-12/23, anti-integrins, and JAK inhibitors-which have been approved for treating UC or CD. Although this wide variety of therapeutics has improved some patients’ lives, they are not universally effective [32]. Moreover, many patients lose responsiveness to therapy over time [33]. Targeting upstream drivers, such as IL-36 cytokines, could have better therapeutic effects, as it would target cytokines downstream, such as TNFα and IFNγ.

## Acknowledgments

This work was supported by NIH Grant 1R01DK120907 (to T.L.D.)

